# Modelling invasive pathogen load from non-destructive sampling data

**DOI:** 10.1101/474817

**Authors:** Nátalia Martínková, Pavel Škrabánek, Jiri Pikula

## Abstract

Where microbes colonizing skin surface may help maintain organism homeostasis, those that invade living skin layers cause disease. In bats, white-nose syndrome is a fungal skin infection that affects animals during hibernation and may lead to mortality in severe cases. Here, we inferred the amount of fungus that had invaded skin tissue of diseased animals. We used simulations to estimate the unobserved disease severity in a non-lethal wing punch biopsy and to relate the simulated pathology to the measured fungal load in paired biopsies. We found that a single white-nose syndrome skin lesion packed with spores and hyphae of the causative agent, *Pseudogymnoascus destructans,* contains 48.93 pg of the pathogen DNA, which amounts to about 1560 *P. destructans* genomes in one skin lesion. Relating the information to the known UV fluorescence in Nearctic and Palearctic bats shows that Nearctic bats carry about 1.7 μg of fungal DNA per cm^2^, whereas Palearctic bats have 0.04 μg cm^−2^ of *P. destructans* DNA. With the information on the fungal load that had invaded the host skin, the researchers can now calculate disease severity as a function of invasive fungal growth using non-destructive UV light transillumination of each bat □s wing membranes. Our results will enable and promote thorough disease severity assessment in protected bat species without the need for extensive animal and laboratory labor sacrifices.

## 1. Introduction

Association of pathogen load with infections transmission stands at the forefront of epidemiological dynamics models, where high pathogen load increases chances of transmission (Wilson et al., 2008). The amount of the pathogen that can infect a new host represents an infectious dose. In natural condition, the infectious dose needs to be transmitted through pathogen shedding via aerosol, direct or vector-mediated oral or bodily fluids exchange. As such, the pathogen is often quantified in sputum, feces, urine or blood, but such measures do not directly reflect the systemic infection or disease severity.

Clinical data support that the overall pathogen load positively correlates with disease severity (Franz et al., 2010; van der Poll and Opal, 2008). Technical and ethical hurdles hinder estimation of the overall load of the pathogen, where the pathogen may occur in different tissues in differing quantities during disease progression (Cunnington, 2015). Thorough sampling can then be possible during autopsy with pathogen quantification in multiple tissue samples, which is sadly late for the patient. Research in infectious diseases aims to prevent lethal outcomes and non-lethal methods must be preferred.

In case of white-nose syndrome (WNS), a fungal infection caused by *Pseudogymnoascus destructans* in hibernating bats (Blehert et al., 2009; Lorch et al., 2011), the disease severity can be estimated from histopathologic examination of wing membrane tissue (Meteyer et al., 2009; Pikula et al., 2017; Reeder et al., 2012). Following skin surface colonization, *P. destructans* invades the skin and forms cupping erosions diagnostic of WNS (Pikula et al., 2017). A cupping erosion is a cup-shaped skin lesion densely packed with fungal hyphae (Meteyer et al., 2009) and the fungus in the cupping erosions produces secondary metabolites that further damage skin of the host (Flieger et al., 2016). Early methodology designed to assess disease severity was lethal, as it required excision and rolling an interdigital segment of the wing in preparation for serial cross-sections used in stained microscopy slides. Number and distribution of the skin lesions were then used to score the severity of the disease (Reeder et al., 2012). More recent methodology utilizes punch biopsy sampling guided by characteristic fluorescence of skin lesions in ultraviolet (UV) light, enabling the bat to survive the examination (Pikula et al., 2017).

Progress from lethal to destructive sampling facilitated examination of bats with higher conservation status for WNS. To further reduce handling and disturbance of bats for research purposes, herein we estimated the fungal load that invaded host tissues in WNS skin lesions. Our approach combines previously published data on fungal load estimated as the amount of *P. destructans* DNA on the wing punch where the wing was without a WNS lesion and fungal load in a wing punch where UV transillumination revealed characteristic fluorescence indicative of WNS lesions. The relationship between the fungal loads measured from the paired biopsies can be used to estimate how much fungal DNA there is in the bat wing tissue, because *P. destructans* fluoresces under UV light after it had formed skin lesions. As the invasive infection progresses, multiple lesions develop and the fungal load in the skin tissue exceeds the fungal load on the wing surface (Martínková et al., 2018). Sampling in the field cannot distinguish between a single microscopic lesion and multiple confluent lesions as both cases manifest as a fluorescing dot when handling a live bat in a cave (Fig. 1). In the laboratory, quantifying pathogen DNA requires enzymatic tissue digestion, meaning that estimating infection intensity with DNA-based methods as well as disease severity with histological microscopy is not possible from the same sample. We therefore simulate possible histopathologic findings in the wing punch digested for pathogen DNA quantification to infer how much *P. destructans* there is in a single WNS skin lesion. We further calculate how the invasive fungal DNA quantity relates to the biology of the infection, and we estimate the number of nuclei in the invading fungal hyphae. We use these results to compare the invasive infection in the Palearctic and Nearctic bats.

**Fig. 1.**
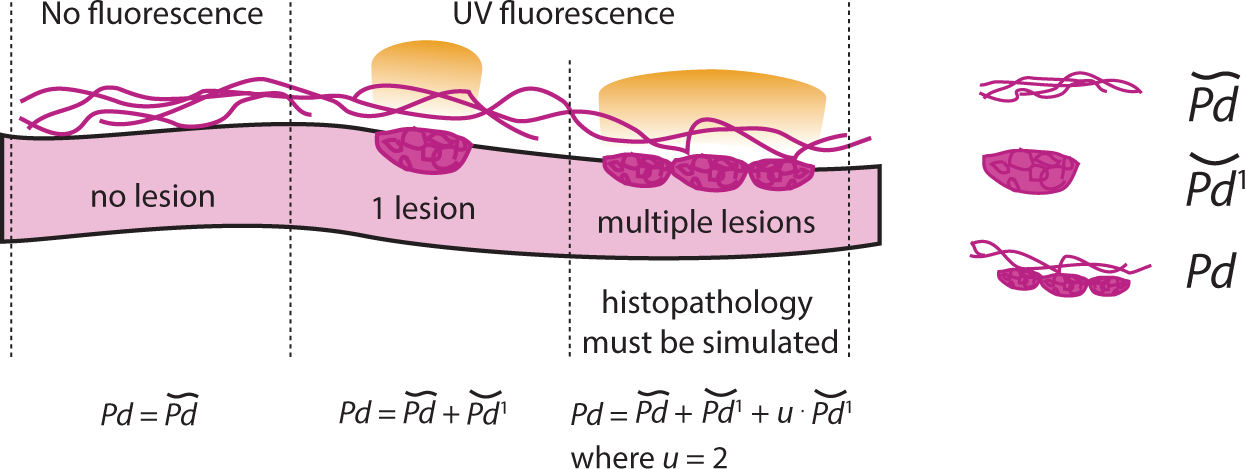
Scheme of the relation between the biological and the mathematical model in estimating fungal load in a skin lesion. Bat wing membrane (pink) was biopsied at two places, once at a wing segment without orange-yellow fluorescence under UV light, and once at a wing segment with a single fluorescing dot. Each biopsy was enzymatically digested and total fungal DNA was quantified with a quantitative polymerase chain reaction. The quantity of fungal DNA, the fungal load, in a biopsy without UV fluorescence represents the fungus that grows on the wing surface (purple lines, 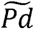). The fungal load in a biopsy with a UV fluorescent dot represents the fungus growing on the wing surface, as well as the fungus that invaded the bat wing and formed cupping erosions diagnostic of white-nose syndrome (purple lines and ovals, *Pd)*. In the field, a single cupping erosion cannot be distinguished from a series of confluent cupping erosions. The unobserved histopathology must thus be simulated to estimate the fungal load in a single cupping erosion 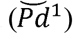.

## 2. Materials and Methods

We used previously published data on fungal load present in a wing punch biopsy (Fig. 1a in Flieger et al., 2016). The data comes from paired wing punch samples from the same bat, where one punch was taken over a membrane segment with an orange yellow fluorescence spot indicating a lesion (Turner et al., 2014) and the other punch from an area where no UV florescent lesions were apparent (Fig. 1). In total, 41 *Myotis myotis* bats with paired wing biopsies were sampled and fungal load as a measure of fungal DNA was quantified with quantitative polymerase chain reaction (Flieger et al., 2016), i.e. sample size *n =* 41.

The dependence of the total fungal load on wing membrane that contains a UV fluorescent lesion diagnostic for WNS *Pd* on the fungal load representing surface colonization on an infected wing membrane 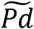 is linear in logarithmic scale (Flieger et al., 2016). The total fungal load in a biopsy with a lesion *Pd* is compounded from the surface skin colonization 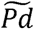 and the invasive fungal growth, where the invasive fungal growth is proportional to the number of skin lesions present in the sample.

The dependence of log_10_(*Pd*) on log_10_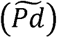 can be used to quantify fungal load in a WNS lesion. Once translated to the quadrant IV of the Cartesian system, the intercept of the linear model α_0_ represents the fungal load in the single lesion 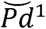. The modified relationship is given as

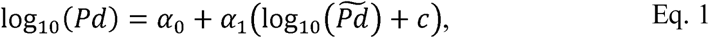

where the constant for translating the data values 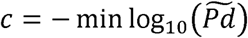. The unknown model parameters *α*_0_ and *α*_1_ can be estimated using a least square method from the set of observations.

Following Eq. 1, the fungal load in a single WNS lesion is 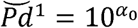 for *α*_1_ ≈ 1. Knowing the fungal load in a single WNS lesion and having enumerated the number of WNS lesions on a wing membrane from their UV fluorescence *n_UV_*, we can calculate the tissue invasive fungal load 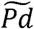 as

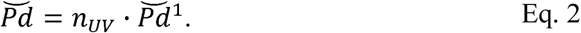

In the empirical study (Flieger et al., 2016), the slope of the regression *α*_1_ = 0.294, meaning that difference in fungal load between a biopsy negative and positive for UV fluorescence is due to additional factors than simple presence of a single WNS lesion in one biopsy. To investigate why there is a departure from 1, we deparsed the covariates confounding total fungal load of the *i*-th single wing membrane biopsy containing a UV fluorescent lesion *pd_i_*. First, the surface of a WNS lesion influences the area, where the fungus colonizes bat wing surface. Second, a UV fluorescent spot observed without magnification in the field might represent a more complex histopathology detectable with microscopy.

### 2.1. Influence of wing surface area of a WNS lesion

The first confounding aspect we considered influences intercept *α*_0_. The surface colonization affects a smaller area in a biopsy with a lesion as the wing surface area of the lesion cannot be considered to contain surface colonization by *P. destructans.* The total fungal load in the *i*-th wing biopsy is

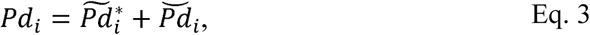

where 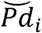 represents fungal load in the UV fluorescent lesion for the *i-*th biopsy, and for this biopsy, 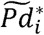 represents a surface colonization satisfying the condition of a sampled area *A*. It holds that

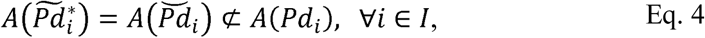

where *I* is a set of sample indices and *I* = {1*,…,n*}.

In the sampling regime used to derive the original model (Flieger et al., 2016), the wing membranes were biopsied with standard 4 mm punch needles. The fungal DNA was thus quantified from the whole wing biopsy with radius *R*, meaning that the surface colonization covers both sides of the biopsy. At least one cupping erosion with radius *r* was present in the biopsy, and 46 % of *M. myotis* biopsies contained multiple cupping erosions (Pikula et al., 2017), i.e. the area covered by a surface colonization in the *i*-th biopsy is given as

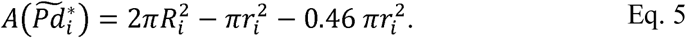

In the empirical studies, the diameter of the *i*-th wing punch biopsy was 2*R_i_* = 4 mm (Flieger et al., 2016) and mean cupping erosion was 2*r_i_* = 86 μm (Zukal et al., 2016) for ∀ *i* ∈ *I*. Solving Eq. 5 numerically shows that 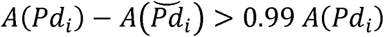 for ∀ *i* ∈ *I*. The effect of presence of the cupping erosion area on the estimation of the fungal load in a WNS lesion is thus negligible.

### 2.2 Influence of unobserved histopathology

The second aspect affects the regression slope *α*_1_ The UV fluorescent lesion recognized without magnification during field sampling could represent a confluent series of cupping erosions (Fig. 1), increasing the relative invasive growth. The fungal load of wing biopsies with UV fluorescence should be revised according to

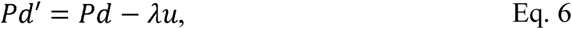

where *Pd′* is the revised fungal load in a wing biopsy with UV fluorescence, *Pd* is the total fungal load estimated from the biopsy, *u* is number of additional cupping erosions, and *λ* is a theoretical fungal load in the UV fluorescent lesion where *λ*∈ [0.1, 10^*α*0^] for *α*_0_ = −1.01527. The value for *α*_0_ was inferred from the least squares regression as per Eq. 1 from the original data, where each biopsy with a UV fluorescence was assumed to contain a single WNS lesion. Note that the published *α*_0_ = 0.015 (Flieger et al., 2016) cannot be used directly, because the original data were not translated following their logarithmic transformation. The published data values are located in the quadrant III of the Cartesian system, where the intercept of the regression does not correspond to the fungal load in a single WNS lesion. In our experience, up to four additional cupping erosions can be associated with one biopsy, i.e. *u* ∈ {0,1,2,3,4} (Fig. 1). Each biopsy in a set of *n* biopsies is associated with one *u* forming a set of alternative histopathologic findings *U* of *n* elements *u*. According to the results (Pikula et al., 2017), probability of multiple UV fluorescent lesions in the set of biopsies is *p*(*u|u* > 0) = 0.46, i.e. *p*(*u|u* = 0) = 0.54. The probability of observing *n_uv+_* additional cupping erosions in biopsies with multiple UV fluorescent lesions is given as *p*(*n_UV+_*) = 0.5 – 0.1*n_UV+_*, where *n_UV+_* ∈ *N_UV+_,* and *N_UV+_ =* {1,2,3,4}. It means that *p*(*u|u* > 0) = 0.46(0.5 – 0.1*u*) for *u* ∈ *N_UV+_*. Given the uncertainty in theoretical fungal load in a UV fluorescent lesion *λ*, there is an infinite number of possible revised fungal loads (Eq. 6) for the available empirical data. We approached the problem using simulations to estimate the effect of multiple UV fluorescent lesions in the biopsy on the regression slope *α*_1_ for sampled combinations of *λ* and *u*. For that purpose, we created a vector ***λ*** with *n_λ_* evenly distributed *λ* where *λ* ∈ [0.1,10^*α*0^], and we included the estimations of the total fungal loads *Pd* for all *n* biopsies with a UV fluorescent lesion into a vector ***Pd***. Further, we generated randomly *n_u_* permutations of the set *U* for each *λ* ∈ ***λ***. The *j*-th permutation of *U* for the *k-*th *λ* is given as ***u**_j,k_* = *ρ_j,k_*(*U*) where *ρ_j,k_* is the *j*-th mapping for the *k*-th *λ*. In total, *n_u_* ⋅ *n_λ_* random vectors ***u*** are generated, where the length of the vector ***u**_j,k_* is *n* for ∀*j*∈{l,…, *n_u_*} and ∀*k*∈{1,…, *n_λ_*}.For the *k*-th *λ* and the *j*-th permutation, the revised fungal loads in wing biopsies are given as

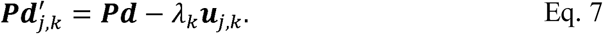

We used the obtained revised fungal loads *Pd′* (Eq. 7) instead of the measured total fungal loads *Pd* while estimating the coefficients *α*_0_ and *α*_1_ of the model Eq. 1 (Fig. 2). Using the least squares estimator, *n_u_* ⋅ *n_λ_* coefficient estimations, 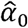 and 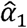, were obtained for the *n_u_ ⋅ nλ* revised datasets. For each *α* ∈ ***λ***, we searched for coefficient estimations 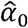 and 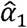 that best match the expected dependence, i.e. we searched for such setting where 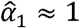. Considering this, we proposed an objective function

**Fig. 2.**
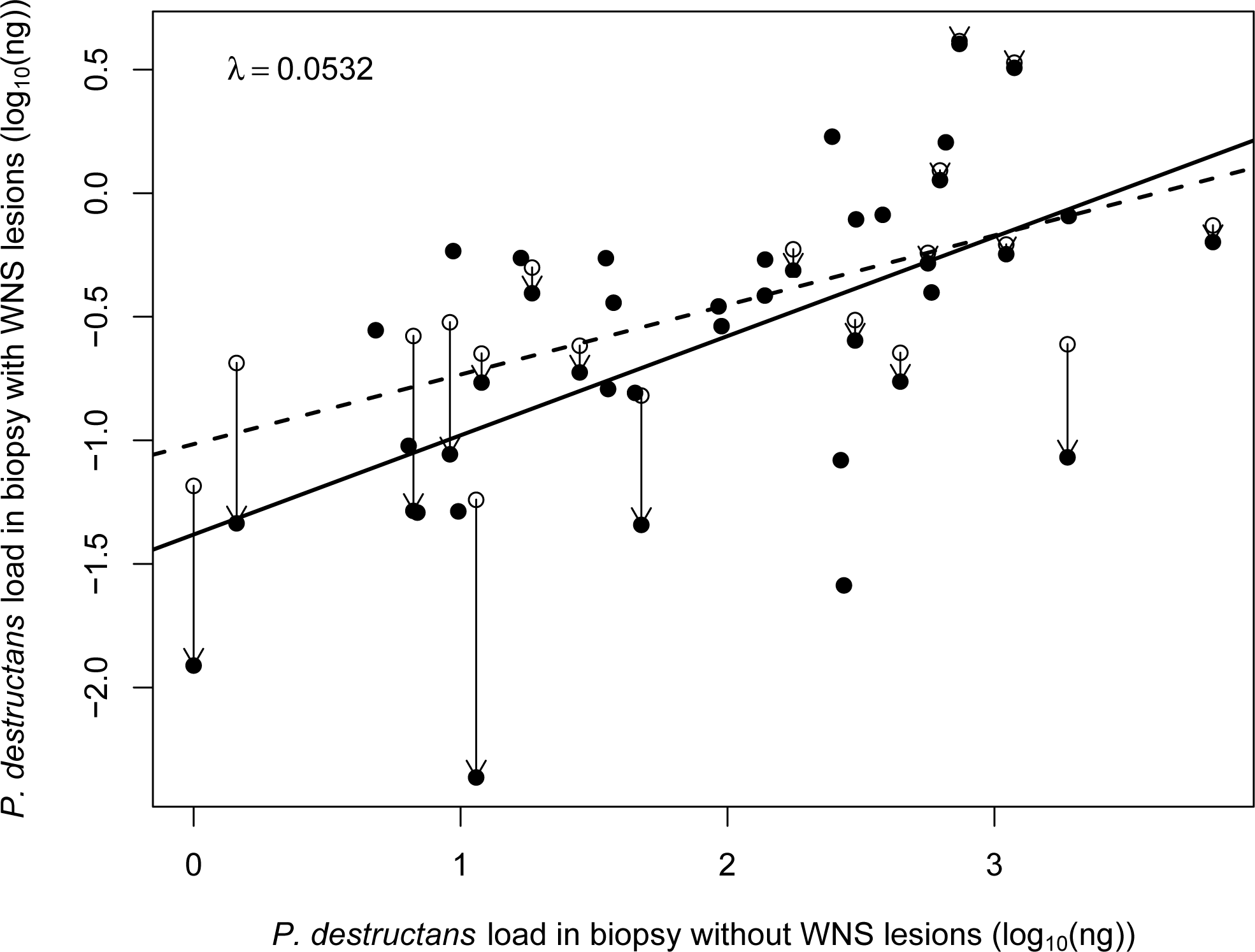
Illustrative modification of fungal load in a bat wing membrane biopsy with UV fluorescence to include simulation of the histopathologic findings. Open circles – measured quantification of *Pseudogymnoascus destructans* DNA in paired biopsies from one bat with the respective regression between fungal load in a wing biopsy without UV fluorescence (and thus without a WNS lesion) and fungal load in a biopsy with UV fluorescence given as a dashed line. Closed circles – adjusted fungal loads, mimicking putative presence of multiple WNS lesions in one biopsy (Eq. 7). The regression line is solid for the adjusted fungal loads. Arrows – direction of change from measured to simulated fungal load per one WNS lesion in the biopsy. Where no arrows are present, the closed circles overlap the measured data in the open circles, meaning that the simulation did not assign more than one WNS lesion in the given biopsy. *λ* – Theoretical fungal load in one WNS lesion as used in the simulation (ng).

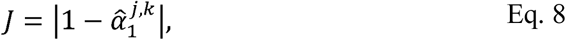

where 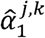 is the estimation of the coefficient *α*_1_ for *λ_k_* and the *j*-th permutation ***u**_j,k_*. For *λ_k_* ∈ ***λ***, the best setting is given as

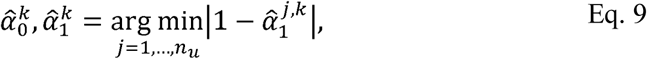

where 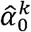 and 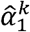 are the best estimates of the coefficients *α*_0_ and *α*_1_ for *λ_k_*. Note that *n_u_* coefficient estimations were obtained for each *λ* ∈ ***λ***, and we calculated fungal load in a WNS lesion as 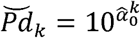.

The study assumes that the first biopsy from the pair represents a wing segment without a WNS lesion, in which *P. destructans* colonized wing surface, and the second biopsy from the pair contains a wing segment with a WNS lesion, where the fungus invaded the tissue. Theoretically, the difference in fungal loads between the two biopsies should be greater than or equal to the fungal load in the WNS lesion. Sampled data that do not conform to the condition are likely influenced by sampling or laboratory artifacts or by as yet unrecognized natural phenomena. To test for the effect of possible unknown bias, we ran a set of simulations on the original and the reduced data. In the case of the reduced data, we used ∀*k* ∈ {1, …, *n_λ_*} those paired biopsies where 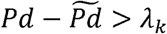.

The equation Eq. 7 allows negative values of revised fungal load in a wing biopsy with a UV fluorescent lesion for some combinations of *λ_k_* and ***u**_j,k_,* which is biologically not feasible. In the absence of *P. destructans,* fungal load must be equal to zero. We considered simulations based on combinations of *λ_k_* and ***u**_j,k_* leading to any *Pd′ <* 0 as failed. Indexes *k* for which any element of 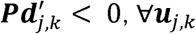 formed a set of failed simulations *D*, which is a subset of all simulations. Fungal load in a single UV fluorescent lesion was estimated from the sampled distribution as

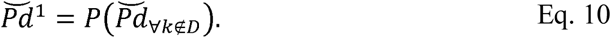

The simulations were run in R (R Core Team, 2018) with custom scripts, implementing equations Eq. 1, Eq. 7, Eq. 8, Eq. 9 and Eq. 10. Data visualization used package *RColorBrewer* (Neuwirth, 2014).

We then used the estimated 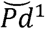 to approximate the number of pathogen nuclei in a skin lesion N. Given the genome size of *P. destructans G =* 30.685 × 10^6^ bp (GCA_000184105.1) and the conversion constant between genome mass and its size *q =* 978 × 1O^6^ (Doležel et al., 2003), the number of *P. destructans* nuclei in a single WNS skin lesion is

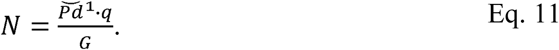

We applied the results of our simulations to previously published data (Pikula et al., 2017) to infer the invasive fungal load in the Nearctic and Palearctic bats using equations Eq. 2 and Eq. 11.

## Results

We ran 1 million simulations of unobserved histopathology during estimation of fungal load in a WNS lesion. For *n_λ_* = 1000, we permuted designation of samples with multiple UV fluorescent lesions *n_u_* = 1000 times for each *λ_k_*. When the theoretical fungal load in a UV lesion *λ_k_* ∈ [0.075,0.097] ng, 127 simulations failed to find any feasible combination of samples with multiple cupping erosions despite attempting one thousand permutations for each *λ_k_* (Fig. 3), indicating that the range might represent an upper limit of fungal load in one WNS lesion in *M. myotis.* Results from 873 successful simulations show that simulating unobserved histopathology improves the objective function value *J* ∈ [0.45,0.80] compared to the result from the original data, where *J =* 0.79 (Flieger *et al.* 2016). The probability density with Gaussian kernel of the simulated fungal load in a lesion has the mean equal to 0.0489 ng and standard deviation equal to 0.01134 ng (Fig. 3). This means that one WNS skin lesion contains 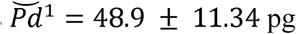 (mean ± SD) of *P. destructans* DNA. The simulated fungal load then translates to 1559 ± 362 pathogen nuclei in a WNS lesion.

**Fig. 3.**
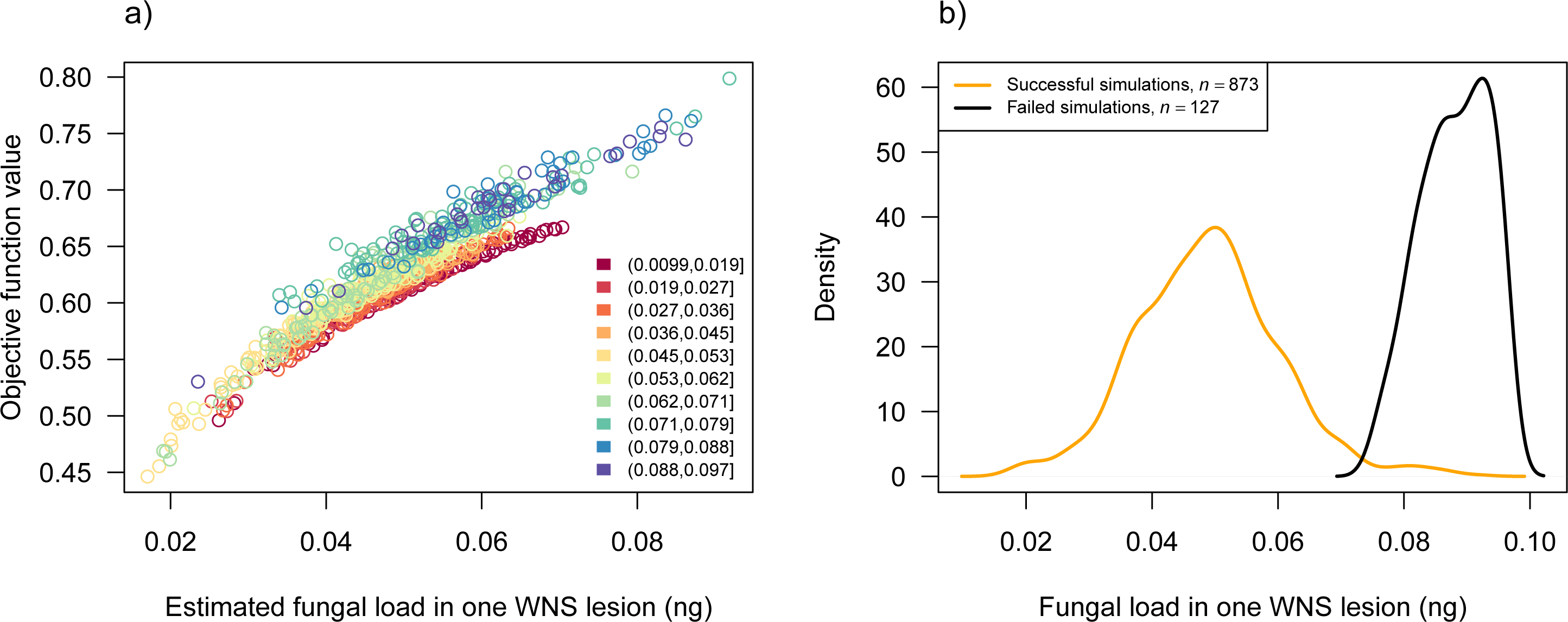
Load of *Pseudogymnoascus destructans* DNA in one UV fluorescent lesion diagnostic for white-nose syndrome in a *Myotis myotis*bat. (**A**) Successful simulations of permuted number of additional WNS skin lesions in a wing biopsy. Theoretical starting values of fungal load in a single WNS lesion are indicated by the colour scheme. (**B**) Density with the Gaussian kernel of the sampled distribution of fungal load in a single WNS lesion in successful (orange) and failed (black) simulations. Simulations were considered failed if all random assignments of additional WNS lesion in a biopsy resulted in at least one adjusted fungal load below zero.

When the data were subset in each simulation to those where 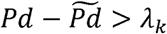, the objective function values further decreased to *J* ∈ [0.28,0.58] and 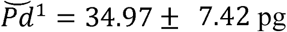 of *P. destructans* DNA. In both simulation modes, using all data and using subsets of data, the theoretical *λ* with the lowest objective function values were about 50.1 to 68.6 pg and the respective estimated 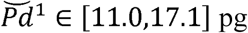.

Using the data on disease severity published by Pikula et al. (2017), the Palearctic bats had *n_uv_ =* 0.78 ± 1.44 WNS lesions per cm^2^ of wing membrane area (*n* = 173), which translates to 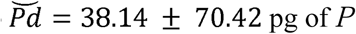 of *P. destructans* DNA or hyphae with 1216 ± 2244 nuclei that invaded the unit area of host tissues. In Nearctic bats with *n_uv_* = 34.73 ± 26.35 UV fluorescent lesions per cm^2^ of wing (*n* = 11), the invasive fungal growth contains 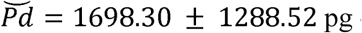 of *P. destructans* DNA, meaning fungal hyphae with 54129 ± 41068 nuclei per cm^2^ of wing area.

## Discussion

The utility in modelling pathogen load from data originating from non-lethal sampling provides unquestionable advantages and insight into disease dynamics. Non-lethal sampling promotes more effective sampling that enables to track distribution, prevalence, spread, infection intensity and disease severity on population level. Our approach simulated unobserved histological severity in a biopsy sample that was used to isolate DNA from the bat as well as the total pathogen biomass in the biopsy. Having established a density distribution of likely fungal loads in one WNS lesion, the researchers can now use the information to infer total fungal biomass that invaded the skin of the hibernating bat. We found that one WNS skin lesion contains about 50 pg of *P. destructans* DNA and thus about 1560 genomic copies in the fungal multinucleic hyphae and spores in the lesion. Translating the value into context of published data on the number of UV fluorescent spots that are indicative of the WNS lesions (Pikula et al., 2017), we found that in some Nearctic bats, 1 cm^2^ of their wing membrane might contain more than 1 ng of pathogen DNA or 54000 pathogen nuclei. The limitation of the present study lies in the fact that values of the objective function Eq. 8 did not approach 0 (Fig. 3a). The objective function minimized difference between the slope of regression from adjusted data with simulated histopathologic findings and the ideal slope equal to 1, when the regression intercept would signify the fungal load in a single WNS lesion (i.e. Eq. 9). The lack of convergence towards the optimum may be due to complexity of WNS pathology in the biopsy we did not consider. The WNS lesions have variable size (Zukal et al., 2016) and some animals develop full thickness invasion where the fungus replaces host tissues across the cross-section of the wing membrane (Pikula et al., 2017). Additional noise in the data is likely introduced with precision of the biopsy punch. Trained personnel stretch a bat wing on a clean, firm surface transilluminated with UV light and circles the target area with a punch needle. Although the punch needles have constant diameter, the sampled wing may differ depending on animal movement, needle slippage or local stretch of the wing membrane. The apparent solution suggests careful sampling where the paired biopsies would be taken from wing area equidistant from joints and bones and the punch site would be chosen with help from magnification to pinpoint single UV fluorescent spots of similar size. At this moment, such data is not available, and our simulation provides the best data-driven approximation of the invasive fungal load during a WNS infection.

We addressed the potential problems in sampling by reducing the dataset to only those observations where the fungal load on a biopsy without a fluorescing WNS lesion was less than fungal load on the paired biopsy by at least the margin of the theoretical fungal load in a single WNS lesion. The change resulted in lower objective function values, but in no simulation was *J* ≈ 0. The uncertainty in the estimate of the fungal load in a single WNS lesion remains influenced by the issues mentioned above. Despite the acknowledged bias, we consider our results useful in infectious disease research of protected bat species, in which using non-destructive methods is warranted. Photography of a bat wing transilluminated with UV light enables enumeration of the WNS lesions in the laboratory and together with estimation of fungal load on the wing surface using a swab sample, these data can be used to calculate infection invasiveness (Martínková et al. 2018). For practical utility in evaluating invasive fungal growth in an infected bat (Eq. 2), we recommend using the fungal load of 49 pg of *P. destructans* DNA in a single WNS lesion. The higher estimate will better incorporate the unconsidered histopathology and also account for presence of confluent WNS lesions that cannot be distinguished on the photograph of the transilluminated bat wing. Using the value of 49 pg of fungal DNA will thus likely better reflect the biological reality of the infection.

Our results provide a valuable tool in assessing invasive infection in endangered hibernating bats on organismal level. Prior to the current study, the disease severity was inferred from focal histopathology in a wing biopsy. Now, the researchers can calculate disease severity as a function of infection invasiveness using non-destructive UV light transillumination (Turner et al., 2014) in conjunction with our results about the fungal load in a WNS lesion.

## Acknowledgement

The study was supported by the Czech Science Foundation (Grant No. 17-20286S). The funding agency had no involvement in study design; in the collection, analysis and interpretation of data; in the writing of the manuscript; and in the decision to submit the article for publication. The authors thank Nancy R. Irwin and Jan Zukal for discussions.

## Contributors

NM and JP conceptualized the study, NM designed and implemented the simulation and collated the data, PŠ formalized the mathematical apparatus, NM and PŠ wrote the manuscript to which all authors contributed.

